# Microeukaryotic predators shape the wastewater microbiome

**DOI:** 10.1101/2023.04.04.535543

**Authors:** Nils Heck, Jule Freudenthal, Kenneth Dumack

**Author notes:** **Correspondence** Kenneth Dumack.

## Abstract

The physicochemical parameters that shape the prokaryotic community composition in wastewater have been extensively studied. In contrast, it is poorly understood whether and how biotic interactions affect the prokaryotic community composition in wastewater. We used metatranscriptomics data from a bioreactor sampled weekly over 14 months to investigate the wastewater microbiome, including often neglected microeukaryotes. Our analysis revealed that while prokaryotes are unaffected by seasonal changes in water temperature, they are impacted by a seasonal, temperature-induced change in the microeukaryotic community. Our findings suggest that selective predation pressure exerted by microeukaryotes is a significant factor shaping the prokaryotic community in wastewater. This study underscores the importance of investigating the entire wastewater microbiome to develop a comprehensive understanding of wastewater treatment.

## 1 Introduction

Prokaryotes are the most abundant microbial entities in wastewater treatment plants (WWTPs), playing a crucial role in the removal of excess nitrogen, phosphorus, and particulate matter (Arregui et al., 2010; Ferrera and Sánchez, 2016; Pan et al., 2018; Wu et al., 2019; Aragaw, 2021). Consequently, extensive research has been conducted to understand the physicochemical parameters that influence the community composition and functioning of prokaryotes in wastewater. Even before the microbial community composition of wastewater was extensively determined, it was recognized that physicochemical parameters affect functioning, and thus wastewater treatment is achieved by altering physicochemical conditions to meet specific processes. For example, denitrification is an anaerobic process, so it functions effectively under anoxic conditions (Lu et al., 2014). In contrast, nitrification is an aerobic process that requires an environment rich in oxygen (Okabe et al., 2011; Ge et al., 2015). Wastewater treatment relies on the coupling of both processes, and thus bioreactors are often periodically aerated. With the advent of molecular tools, it was discovered that physicochemical parameters also significantly influence the community composition of prokaryotes. For instance, pH is known to correlate with the relative abundance of certain prokaryotic taxa, and seasonal effects are frequently reported (Gao et al., 2016; Herold et al., 2020; LaMartina et al., 2021). Season, however, is a sum of numerous environmental changes, for instance, in the light regime and temperature. Accordingly, seasonality affects community composition and subsequently function and performance in wastewater treatment through accompanying environmental changes whose interplay is often poorly understood (Kim et al., 2013; Liu et al., 2016; González-Camejo et al., 2018; Schages et al., 2020).

Apart from prokaryotes, the microbiome in WWTPs also includes microscopic animals and microeukaryotes such as fungi and protists. Although microscopic animals, fungi, and phototrophic protists (i.e., algae) are occasionally included in surveys of wastewater, predatory protists were long marginalized (Cuellar-Bermudez et al., 2017; Freudenthal et al., 2022). This knowledge gap is mainly due to the challenges in taxonomic identification and enumeration. While primer-based metabarcoding is currently the most commonly used method to assess microbiomes, protists are genetically diverse and paraphyletic, leading to contradictory results in protist assessments (Lentendu et al., 2014; Sibbald and Archibald, 2017; Hirakata et al., 2019; Maritz et al., 2019; Burki et al., 2020). However, with shotgun meta-omics, it is now possible to assess the protistan community without any primer bias (Freudenthal et al., 2022). Furthermore, these techniques enable the assessment of the entire microbiome, from prokaryotes to microbial eukaryotes. Nevertheless, these promising methods have rarely been used to investigate microbial communities in wastewater, and if so, the data were mostly not screened for protists.

Despite past methodological hurdles, our knowledge of how microbial eukaryotes, particularly predators, affect wastewater treatment still relies largely on decades-old surveys or experiments (Foissner, 2016). While these findings can describe certain outcomes, they are often insufficient to link these changes to respective species in the microbial community. Predatory protists were found to be involved in floc formation, clarification, and the removal of parasites, but whether their specific predation pressure significantly shapes the prokaryotic community composition in wastewater is unknown (Lee and Welander, 1996a, 1996b; Pauli et al., 2001; Arregui et al., 2010; Freudenthal et al., 2022).

This study aims to address this knowledge gap by analyzing a publicly available metatranscriptomic dataset provided by Herold et al. (2020), who sampled the anoxic tank of a WWTP every week for about 14 months. The dataset allowed us to identify the abiotic and biotic factors that change the community composition of the entire wastewater microbiome over time. Our specific objective was to assess the wastewater microbiome in its entirety, identify which environmental factors shape its composition, and particularly investigate whether biotic interactions shape the community.

## 2 Material and Methods

### 2.1 Data access, quality filtering, and assembly

We utilized a publicly available dataset provided by Herold et al. (2020) for this study. Briefly, the samples were collected on a weekly basis over a period of 14 months from the anoxic (denitrification) zone of the aeration tank of a municipal biological wastewater treatment plant located in Schifflange, Luxembourg (49°30⍰48.29⍰ ⍰N; 6° 1⍰4.53⍰ ⍰E; Herold et al., 2020). The treatment plant in Schifflange, after primary clarification, uses an activated sludge process. This takes place in two aeration basins, which have an outer aerobic nitrification zone and an inner anaerobic denitrification zone each, with a total volume of 17400 m^3^ (SIVEC last accessed 6.21.23). Samples were taken from floating sludge islets in the denitrification zone. After the samples had been collected, RNA was extracted as part of a sequential co-isolation procedure. For RNA extraction and cDNA library preparation, please refer to Herold et al. (2020) and Roume et al. (2013). The code for the presented analyses including set parameters is available on GitHub under the following link: https://github.com/N-Heck-1/Microeukaryotic-predators-shape-the-wastewater-microbiome. The quality of the raw data received was assessed using fastqc v. 0.11.9 (Andrews, 2010). We used TrimGalore v. 0.6.7 (Krueger et al., 2021) to detect and remove adapters and perform quality trimming, which removed bases with a quality score of <30 and the last ten bases of the 3’ end. Mothur v. 1.45.3 (Schloss et al., 2009) was used to assemble the paired-end reads into contigs. The contigs were screened for a minimum length of 90 bp and a maximum of 2 bp in ambiguities and mismatches after determining appropriate screening parameters.

### 2.2 Taxonomic assignment

After quality filtering, we screened the sequences using the BLASTN algorithm (Camacho et al., 2009) against the respective databases. To identify eukaryotic taxa, we used the PR v. 4.11.1 database (Guillou et al., 2013), and for prokaryotic taxa, we used the SILVA 138 SSU Ref Nr. 99 database (Pruesse et al., 2007). We kept only the best hit for each result, and filtered the results using an e-value threshold of 1e^-35^ for PR^2^ and 1e^-30^ for SILVA results, a bit-score threshold of 140, and an identity threshold of >92% across the data set. As our investigation focused on the microbial community, we removed sequences derived from chloroplasts, macroscopic Metazoa, and Embryophyta. Taxon counts were binned at the genus level to form the OTUs for all further analyses. We performed data filtering and analysis in R v. 4.0.5 (R Core Team, 2021), using the packages lattice v. 0.20-41 (Deepayan, 2008), tidyr v. 1.1.3 (Wickham and Girlich, 2022), dplyr v. 1.0.7 (Wickham et al., 2022), ape v. 5.6-2 (Paradis, 2006), and vegan v. 2.5-7 (Dixon, 2003; Oksanen et al., 2020). Additionally, we used ggplot2 v. 3.3.4 for visualization (Wickham, 2016).

### 2.3 Removal of outliers

To ensure the uniformity of sequencing results, we initially screened the data and removed two samples with much lower sequencing depth and which were sampled much earlier than the others (i.e., 2010-10-04, 2011-01-25). We also removed singleton to quintuplet OTUs before proceeding to further analysis. To identify potential outliers among samples, we utilized multivariate dispersion based on normalized and Bray-Curtis transformed data (Bray and Curtis, 1957) and generated an NMDS plot using the metaMDS function from the vegan package. Based on the plot, we excluded three additional samples (i.e., 2011-04-05, 2011-03-29, 2011-03-21; see Supplementary Figure 1). We then produced rarefaction curves to assess sequencing depth across the remaining samples and excluded those with a depth below 1e^-6^ ribosomal reads (i.e., 2011-11-23, 201-11-16, 2011-11-29; see Supplementary Figure 2). To facilitate comparability between samples, we rarefied the data to a depth of 1,066,681 ribosomal reads per sample, resulting in the removal of 43 rare OTUs out of a total of 4386.

### 2.4 Statistical analyses

To visualize temporal changes in community composition, stacked bar plots were generated. An NMDS plot was also computed to visualize the impact of environmental factors on the prokaryotic and eukaryotic communities. The environmental variables considered were season, water temperature, conductivity, and oxygen saturation, which were measured as described in Herold et al. (2020). The potential effect of these parameters on community composition was explored using the envfit and orditorp functions from the vegan package. To investigate the influence of biotic factors on prokaryotic and eukaryotic community structure, principal coordinate analysis (PCoA) was employed to compute a measure of community structure using the pcoa function from the ape package (Gower, 1966). The PCoA revealed that the first two axes explained 67.7% and 12.9% of the variation in the prokaryotic community, respectively, and 41.5% and 17.8% of the variation in the microbial eukaryotes. The coordinates of the axes with the highest explanatory power were used for further analyses. To investigate the impact of environmental variables and biotic interactions on the eukaryotic and prokaryotic communities, PERMANOVA was performed using the adonis function from the vegan package (Anderson, 2001). To investigate the factors and interactions that contribute significantly to the variation in prokaryotic and eukaryotic communities, we employed variance partitioning analysis. To address the issue of numerous zero values in the abundance data, a Hellinger transformation was applied prior to analysis using the varpart function from the vegan package (Legendre, 2008). To identify the variables that significantly explained the variation in community composition, we conducted forward selection using the ordistep function from the vegan package.

### 2.5 Network analyses

To visualize and locate biotic interactions, networks were calculated. Sample heterogeneity caused by rare taxa was reduced to lessen their influence on network precision through co-absence (Röttjers and Faust, 2018). A prevalence filter was applied, excluding all OTUs present in less than 50% of the samples, resulting in 1559 OTUs being included in the network analyses. FlashWeave v. 0.19.0 (Tackmann et al., 2019) was chosen to investigate the multitude of complex associations within the WWTP microbiome. As a cross-sectional tool for network inference, it is well-suited for data with varying time intervals (Gerber, 2014). FlashWeave provides good scalability for large data sets, performs well on data with many heterogeneous samples, and enables the inclusion of environmental data (Matchado et al., 2021). The tool was run in sensitive mode with default settings in the Julia v. 1.7.3 environment (Bezanson et al., 2017). The resulting network was simplified for visualization in Cytoscape v. 3.9.1 (Shannon et al., 2003), by aggregating calculated nodes at the order level. This step included the influences of unique OTUs as edges in the network. If edges with conflicting signs were introduced by this step, both were kept, reflecting evenly contrasting interactions (Röttjers and Faust, 2018).

On the OTU level, 2673 edges were found between 1530 nodes and aggregated to 1767 edges between 327 nodes at the order level. Associations within functional groups (here defined as Metazoa, fungi, bacteria, Archaea, and protists) accounted for 883 edges. Among these, 610 edges represented associations among protists, 228 edges among bacteria, and 51 among fungi. Node size was chosen to denote the total number of ribosomal reads across all samples for that order, and node color indicated the number of OTUs that were aggregated. Edge thickness was related to the number of individual edges between the two orders, while edge color denoted the sign. To further ease the visualization of the network, only edges between functional groups and their respective nodes were shown, and nodes were grouped by higher taxonomic levels.

## 3 Results

The microbial community in the wastewater treatment plant underwent seasonal changes, with the eukaryotic community experiencing more pronounced changes compared to the prokaryotic community (Fig. 1). Prokaryotic seasonal clusters were not clearly separated (Fig. 1A), while a separation of gradually changing winter and summer clusters was observed for eukaryotes (Fig. 1B).

**Figure 1:**
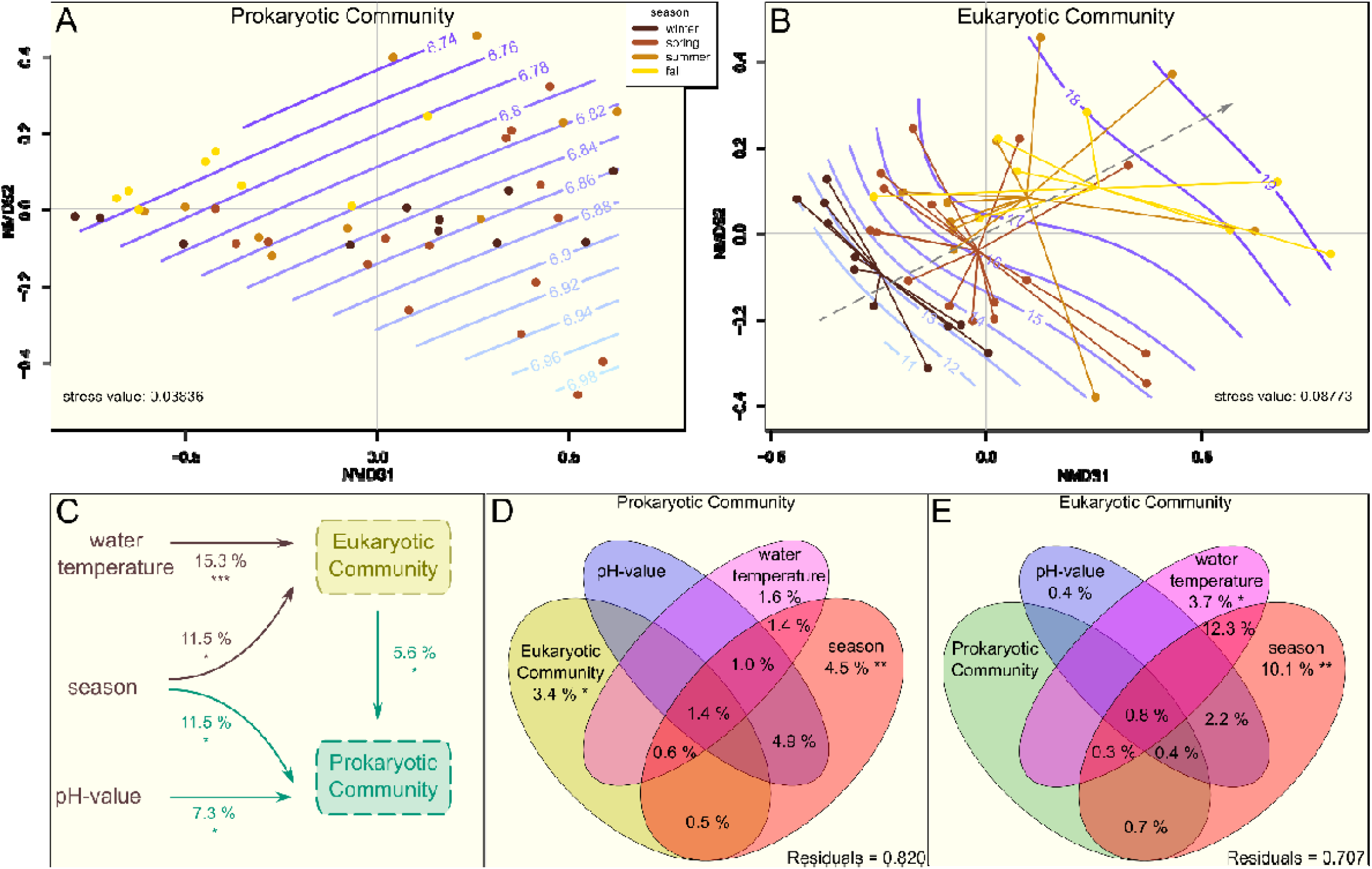
Influences on the prokaryotic and eukaryotic community composition. **(A)**The prokaryotic wastewater community is depicted, with the strongest abiotic shaping force, pH, highlighted – seasonal effects are not clearly visible. In contrast, the eukaryotic community composition is shaped by season and related water temperature **(B)**. The color of each point indicates the season (A+B). Smooth surfaces (blue lines) represent the influence of the abiotic parameter with the highest explanatory power on the respective community compositions, i.e., pH for prokaryotes and water temperature for eukaryotes, respectively. The central arrow in B indicates the orientation of seasonal cluster centromeres along the temperature gradient. **(C)** Predictors for prokaryotic and eukaryotic community composition are shown. Arrows indicate significant influences on the microeukaryotic and prokaryotic communities, respectively. The structure (PCoA1) of the eukaryotic community was used as a predictor for prokaryotic data, and the structure (PCoA1) of the prokaryotic community was used as a predictor for eukaryotic data. The R^2^-values indicate the percentage of the respective factor explained. Asterisks indicate the significance codes: * for 0.05 ≥ p > 0.01, ** for 0.01 ≥ p > 0.001, and *** for p ≤0.001. Only significant factors are shown. **(D+E)** Venn diagrams show the partitioned variance explaining the composition of the prokaryotic **(D)** community and the eukaryotic community. **(E)**. Significant effects according to forward selection are marked by stars: 0.05 ≥ * > 0.01 ≥ **. Note that here too a significant effect of the eukaryotic community composition on the prokaryotic community composition was found **(D)** and that a large percentage of the seasonal effect explaining the variance of the eukaryotic community composition overlaps with a temperature effect **(E)**.

Seasonal changes shaping the microeukaryotic community can be traced back to a response of the microeukaryotes to changes in season-dependent water temperature (Fig. 1 C). This observation was supported by variance partitioning, which revealed that both season and temperature accounted for significant proportions of the variation in the microeukaryotic data, with a substantial overlap of 12.3% (Fig. 1 E). In contrast, we did not observe a clear seasonal response of the prokaryotic community to changes in water temperature. To determine which seasonal changes, if not water temperature, influenced the changes in the prokaryotic community, it was tested whether the seasonal variation of the prokaryotes could be explained by the changes found in the eukaryotic community composition. The analysis revealed that the variation in the prokaryotic community composition (Fig. 1 C) was indeed explained by the variation in the eukaryotic community composition (PCoA1). Supporting these findings, approximately 14% of the variation in the prokaryotic dataset was attributed to seasonal changes, while a smaller but still significant proportion (about 6%) was explained by the eukaryotic community composition, as determined by variance partitioning (Fig. 1 D). In addition to the described seasonal and biotic effects, the pH value was found to explain variation in prokaryotes, too (Fig. 1 A, C). The other tested chemical parameters showed no effects.

Since we found a shaping influence of the eukaryotic community composition on the prokaryotic community composition, we investigated which pro- and eukaryotic taxa changed over time. We found a total of 4343 operational taxonomic units (OTUs) composed of 47,995,763 SSU rRNA sequences in the bioreactor. In general, the prokaryotic community was dominated by the orders Enterobacterales (Gammaproteobacteria), followed by Nitrososphaerales (Crenarchaeota), Methanosarciniales (Halobacterota), and Bacillales and Lactobacillales (Firmicutes; Fig. 2 A). The eukaryotic community was dominated by Vannellida (Amoebozoa), making up almost half of all eukaryotic rRNA reads, followed by the second most abundant order Peritrichia (Ciliophora). Among the other highly abundant groups were Prostomatea (Ciliophora), Kinetoplastida (Discoba), Cryomonadida (Cercozoa), Trebouxiophyceae (Chlorophyta, green algae), and the fungal taxa Saccharomycotina and Pezizomycotina (Fig. 2 B).

**Figure 2:**
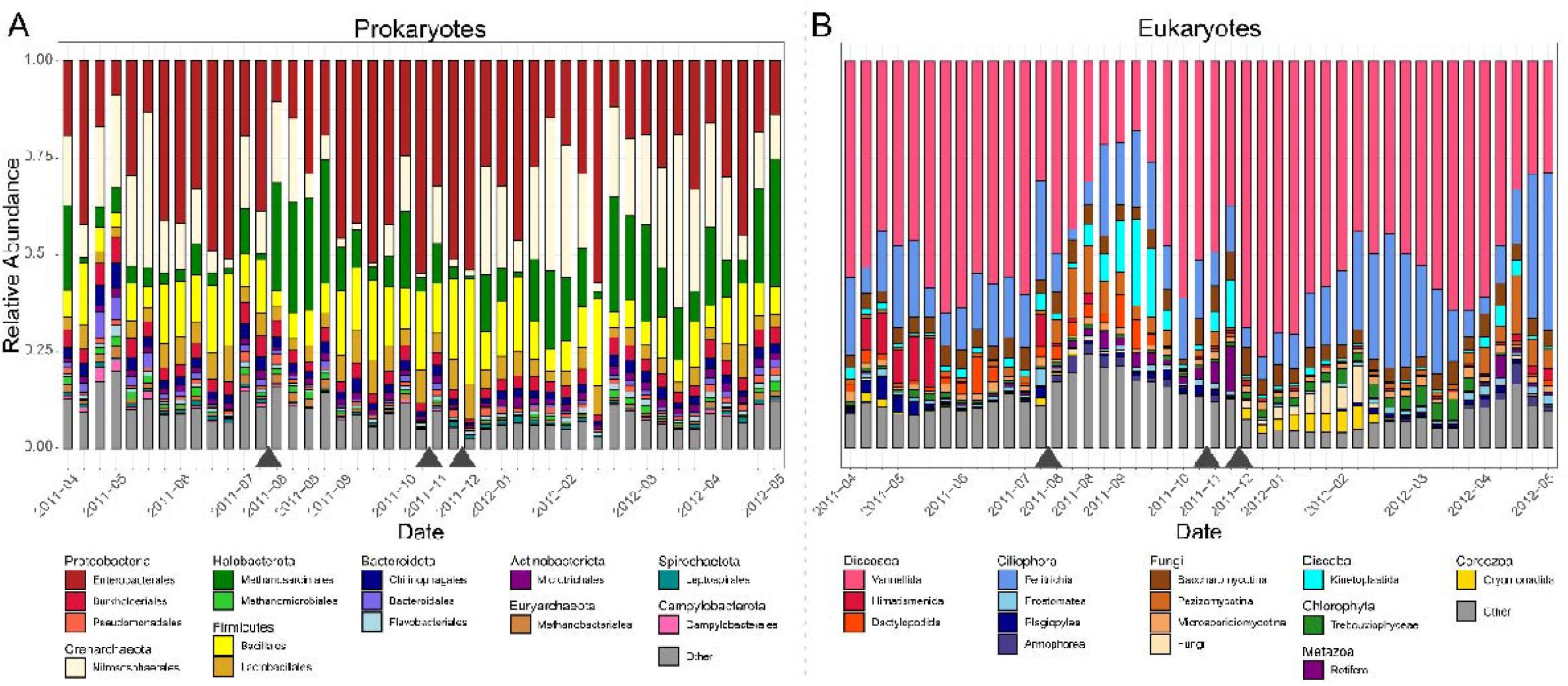
Microbial community composition in the anoxic compartment of a wastewater treatment plant over 13 months. Stacked bar plots display the relative abundance of the 15 most abundant prokaryotic **(A)** and eukaryotic **(B)** orders. Each bar represents the community in one sample, and gray triangles on the x-axis indicate gaps in the dataset due to the removal of low-quality samples.

Many of the dominant eukaryotic orders displayed a higher relative abundance in samples taken at lower water temperatures in winter and early spring, such as Vannellida (Amoebozoa), Cryomonadida (Cercozoa), and the green algae Trebouxiophyceae (Chlorophyta; Supplementary Figure 3A-C). In contrast, Himatismenida and Dactylopodida (both Amoebozoa; Supplementary Figure 3D), as well as Pezizomycotina (fungi) showed an increase in their relative abundance with higher water temperature, mostly from April to September (Fig. 2B). Kinetoplastida (Discoba) and Rotifera also showed higher abundances from late summer to early winter, while both were hardly detectable from December to April.

To investigate which biotic interactions in the microbial community may have led to the observed changes, we employed network analysis. The most pronounced result was the numerous negative associations between protists and bacteria (∼80.1%), potentially indicating predator-prey relationships. A total of 785 edges indicated associations between functional groups, with 407 (∼51.8%) between fungi and protists, 159 (∼20.3%) between bacteria and protists, and 77 (∼9.8%) between bacteria and fungi (Fig. 3). Most associations between protists and fungi were positive (∼66.9%).

**Figure 3:**
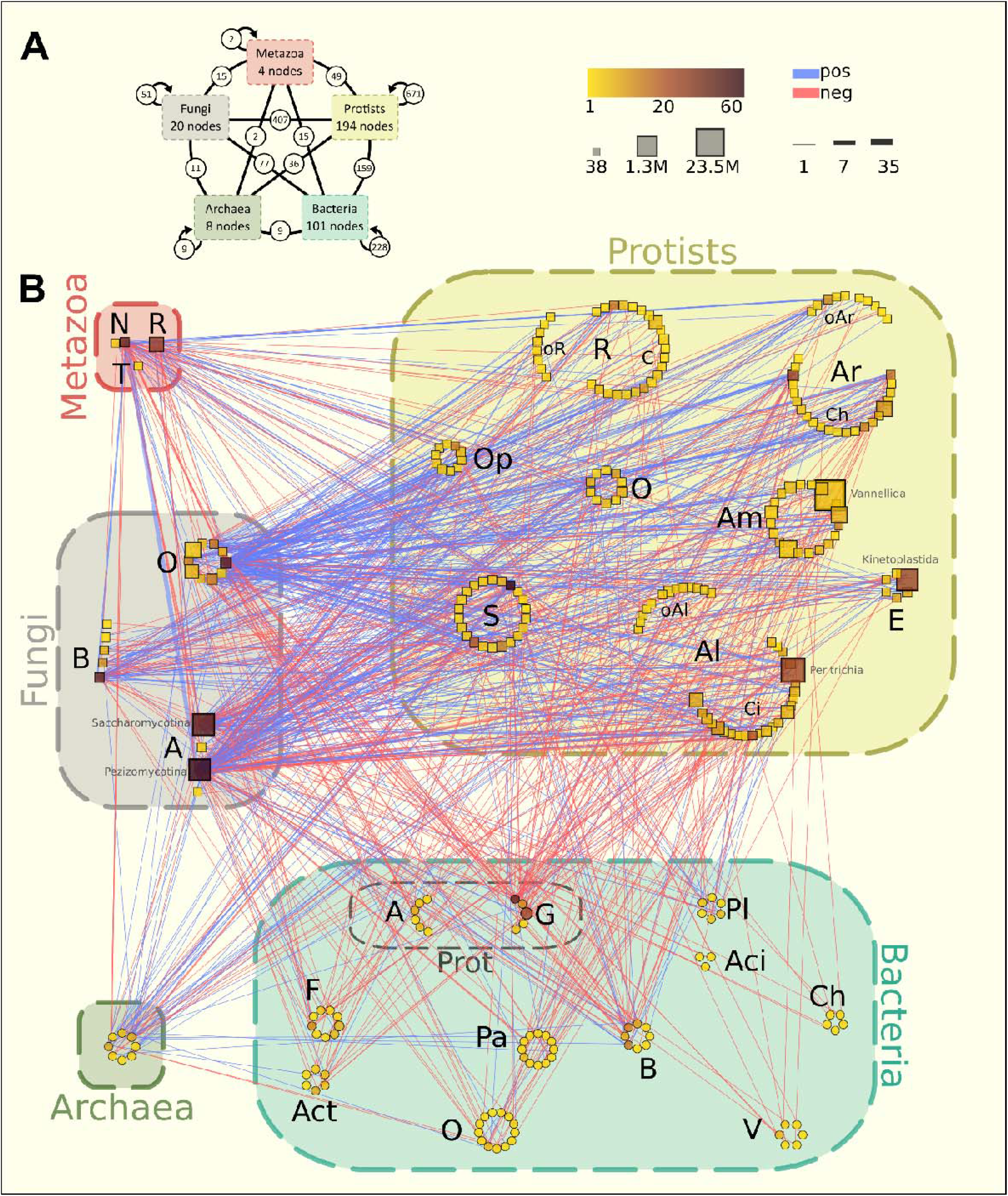
Co-occurrence network of microbial orders in the anoxic compartment of a biological wastewater treatment plant. Nodes were aggregated at order level. **(A)** shows a summary of the network presented in **(B)**, highlighting the number of nodes in each functional group and the number of edges within and between each group. In **(B)**, the node colour indicates the number of genera included and node size the total number of reads over all samples, as shown in the legend. Blue edges show positive association, red edges show negative association, and the thickness of the edges shows the number of genera involved. Only edges between the functional groups are displayed in the network itself **(B)**, and only nodes with at least one edge are displayed. Abbreviations for Metazoa: N=Nematoda, R=Rotifera, T=Tardigrada. Abbreviations for Fungi: A=Ascomycota, B=Basidiomycota, O=Other. Abbreviations for Protists: Al=Alveolata, Am=Amoebozoa, Ar=Archaeplastida, C=Cercozoa, Ci=Ciliophora, Ch=Chlorophyta, E=Excavata, O=Other, oAl=other Alveolates, oAr=other Archaeplastida, Op=protistan Opisthokonta, oR=other Rhizaria, R=Rhizaria, S=Stramenopiles. Abbreviations for Bacteria: A=Alphaproteobacteria, Aci=Acidobacteriota, Act=Actinobacteriota, B=Bacteroidota, Ch=Chloroflexi, F=Firmicutes, G=Gammaproteobacteria, O=Other, Pa=Patescibacteria, Pl=Planctomycetota, Prot=Proteobacteria, V=Verrucomicrobiota.

## 4 Discussion

The performance of a wastewater treatment plant is highly dependent on the composition and activity of its microbial community (Arregui et al., 2010; Liu et al., 2016; Aghalari et al., 2020). Therefore, it is crucial to have a detailed understanding of the factors influencing the microbial community to ensure consistent and efficient WWTP performance. This study reveals that the seasonally changing microeukaryotic community composition is a significant factor shaping the prokaryotic community.

Researchers have been investigating for decades the environmental parameters that shape the prokaryotic community composition in WWTPs (Kim et al., 2013; Liu et al., 2016; Herold et al., 2020). Although the potential importance of eukaryotes for wastewater treatment has been described, it has not received much attention due to a lack of suitable methodology (Pauli et al., 2001; Arregui et al., 2010; Foissner, 2016). Accordingly, a previous study reported no clear seasonality for protists (Utz and Bohrer-Morel, 2008), while another study found that seasonally changing water temperature influenced protistan growth (Hirakata et al., 2019). In this study, we found clear seasonal variations in both prokaryotes and eukaryotes. Although the temperature in the anoxic tank fluctuated seasonally with the ambient temperature, the seasonal effect on the prokaryotic community composition could not be explained by a direct effect of water temperature. Instead, we conclude that the eukaryotic community composition was impacted by seasonally changing water temperatures, which, in turn, shaped the composition of the prokaryotic community. This finding underscores the importance of focusing on the whole microbial community and its interactions to facilitate our understanding of wastewater treatment processes.

For the first time in a WWTP, we were able to disentangle a shaping force exerted by the microeukaryotic community on the prokaryotic community composition. Our use of network analysis revealed numerous negative associations between eukaryotic and prokaryotic groups, further visualizing the shaping impact that had been found statistically. Such negative edges may have diverse ecological explanations, including competition and predation (Faust and Raes, 2012). We argue that this shaping force is largely caused by protistan predation. All of the most numerous protists in this study are known bacterivores (Böhme et al., 2009; Vaerewijck et al., 2011; Samba-Louaka et al., 2019), including sessile peritrich ciliates like Vorticella, which are considered important for treatment efficiency, effluent clarity, and nitrification, as well as Kinetoplastida and Cryomonadida, both of which are known to feed on wastewater bacteria (Pauli et al., 2001; Arregui et al., 2012; Foissner, 2016; Liu et al., 2016; Pohl et al., 2021; Freudenthal et al., 2022). The most abundant protistan taxon in this study, bacterivorous Vannelida, is often absent from wastewater surveys that rely on primer-based methods, but studies based on culturing or metatranscriptomic methods have repeatedly reported their presence in WWTP bioreactors (Ramirez et al., 2015; Freudenthal et al., 2022). We thus consider this to be a primer bias against the amoebozoan Vannellida (Urich et al., 2008; Geisen et al., 2015; Fiore-Donno et al., 2016).

It is still uncertain to what degree changes in prokaryotic community composition caused by protistan predation affect their functioning. It is known that there is a certain redundancy in prokaryotic functioning in wastewater, where multiple different prokaryotes are involved in the same nitrogen removal processes (Pan et al., 2018). Given that protists feed selectively, it is reasonable to assume that not all of these taxa are preyed upon at the same rate. Such redundancy suggests that changes in the prokaryotic community due to protists may only have a limited effect on wastewater treatment functions, but this requires further investigation (Ju et al., 2014). Our analysis revealed a high abundance of Nitrosphaerales, which are ammonia-oxidizing Archaea involved in wastewater nitrification with increased abundance in hypoxic conditions (Park et al., 2006; Limpiyakorn et al., 2013; Rodríguez et al., 2015; Ferrera and Sánchez, 2016). However, it is not well understood whether, and at what rate, protists prey on Archaea.

In addition to predation, protists also compete with prokaryotes. For example, phototrophic protists (i.e., algae) are known to compete with prokaryotes for nutrients such as nitrogen and phosphorus (Cuellar-Bermudez et al., 2017; González-Camejo et al., 2018). Nevertheless, we argue that competition for nutrients is likely less significant than predation throughout the year. Phototrophic protists were generally less abundant than predatory ones, but they peaked in their abundance during winter. Therefore, eukaryote-prokaryote competition may be more pronounced in winter.

## 5 Conclusion

The results presented in this study, along with several other recent studies, emphasize the importance of giving more attention to microeukaryotes, particularly protists, in the investigation and improvement of wastewater treatment (Foissner, 2016; Assress et al., 2019; Freudenthal et al., 2022). Additionally, we demonstrate the significant benefits of primer-independent metatranscriptomics in generating comprehensive datasets that encompass the entire microbial diversity.

## 6 Acknowledgements

We thank Marcel Dominik Solbach for providing the drawing of the wastewater treatment plant that we included in the graphical abstract.

## Notes

### Competing Interest Statement

The authors have declared no competing interest.

### Summary of Updates

We followed reviewer's comments

