## Supplementary files for "Microeukaryotic predators shape the wastewater microbiome"

Corresponding author Kenneth Dumack

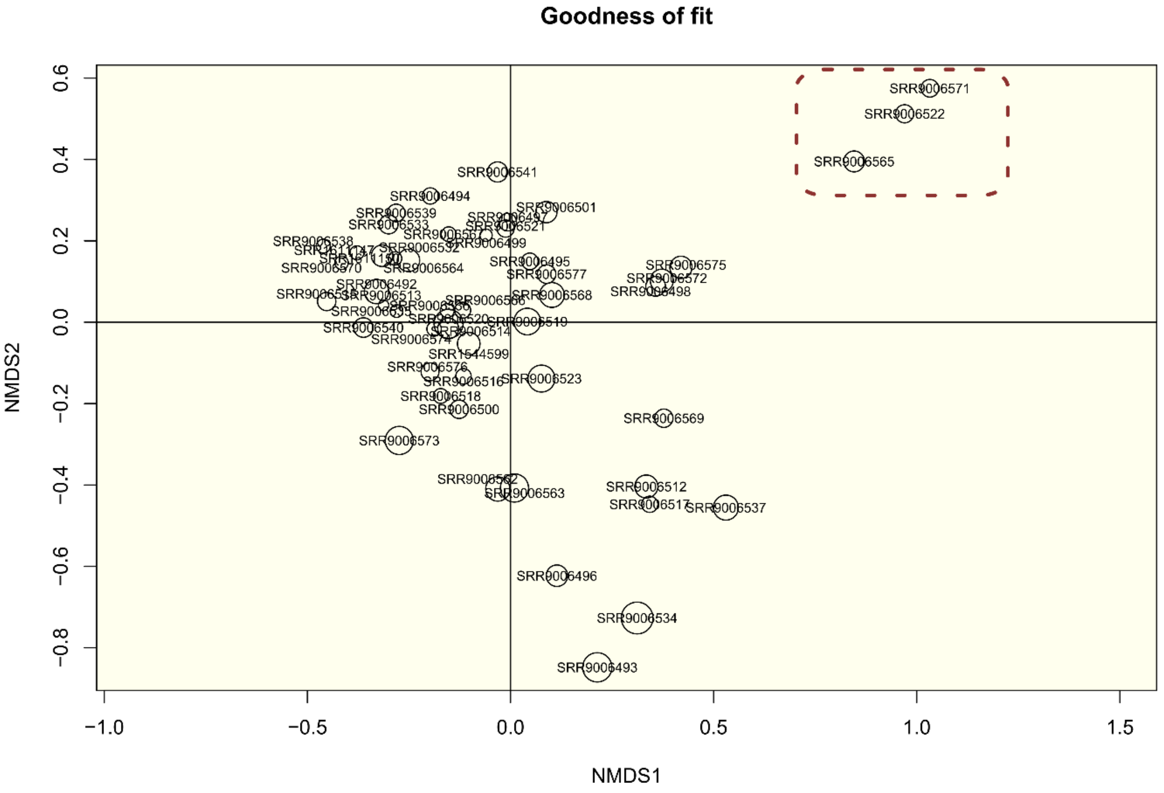


**Supplementary Figure 1: Nonmetric multidimensional scaling plot used to determine outliers.** Each site represents one sample. Circle size represents the goodness of fit. Outlier samples to be removed are marked by a red frame.

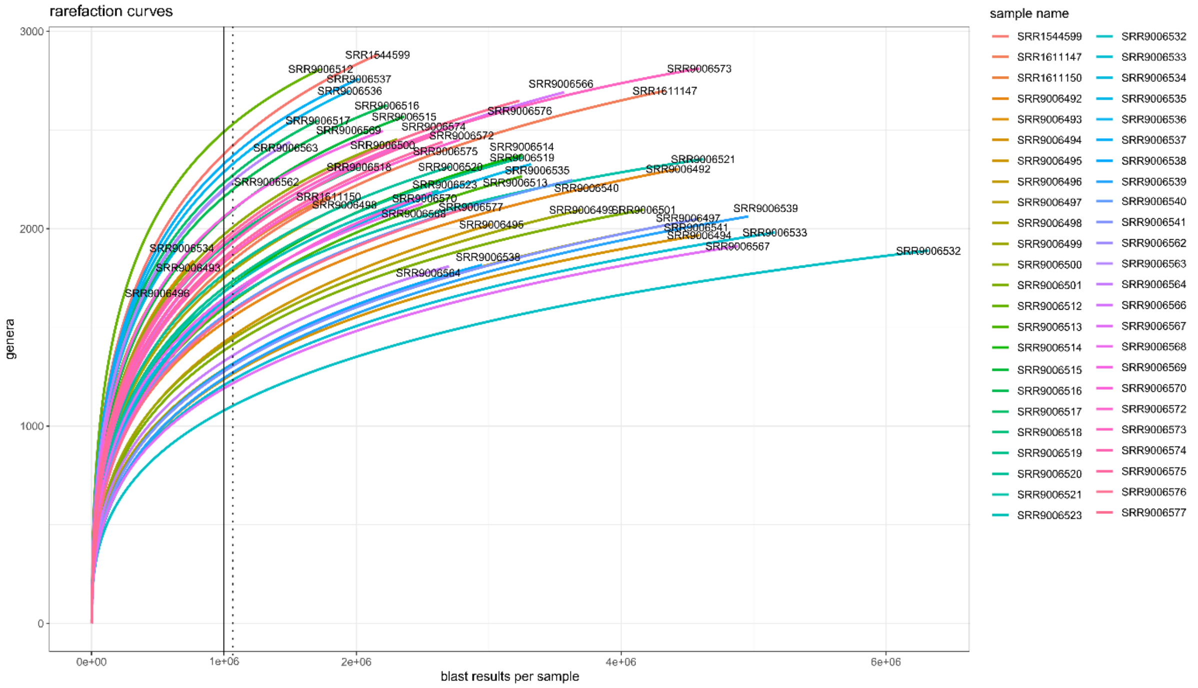


**Supplementary Figure 2: Rarefaction curves.** The number of OTUs at the genus level is plotted against the number of reads for each sample. The black line denotes $1e^6$ rRNA reads, which was chosen as the cutoff for sequencing depth. The dotted line is set at the number of reads to which all samples were rarefied – 1,066,681.

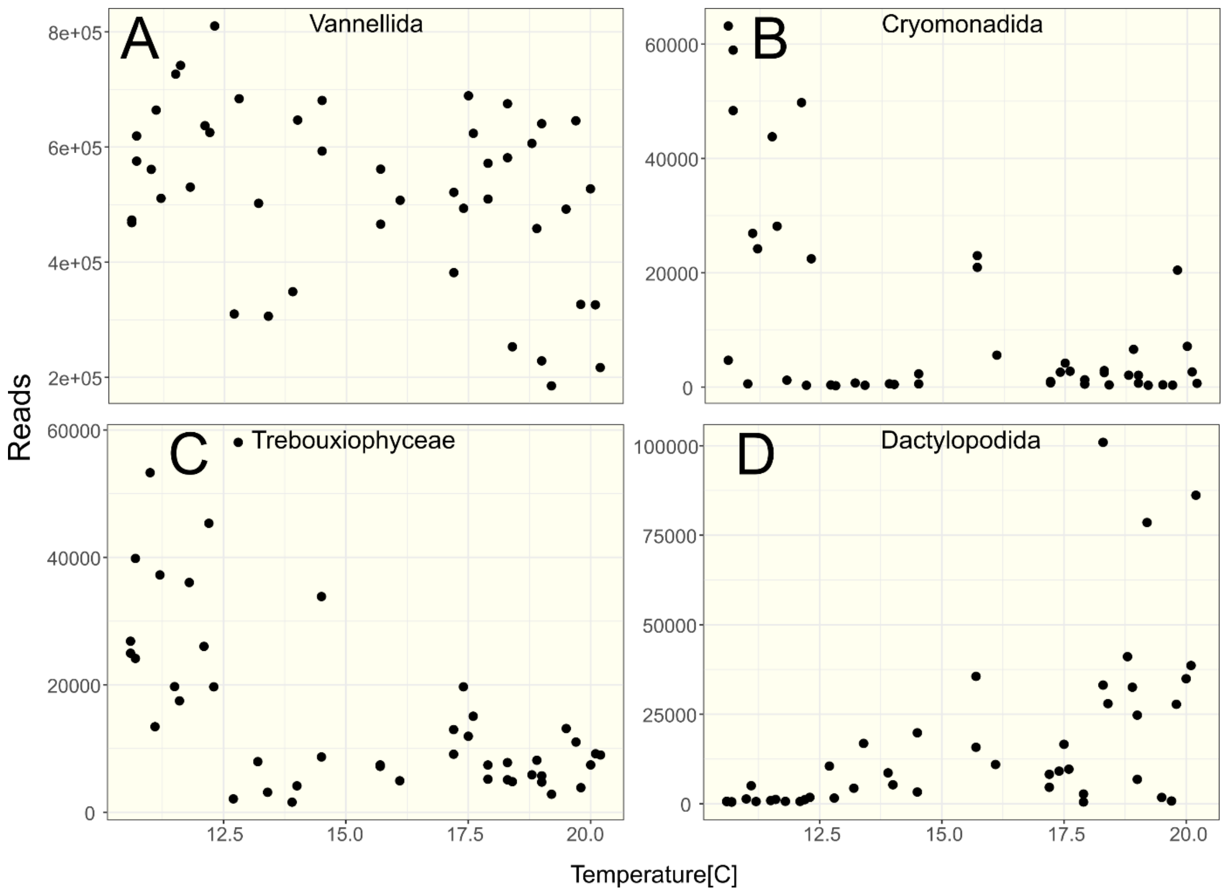


**Supplementary Figure 3: Influence of temperature on selected prominent eukaryotic orders.** The abundance of selected eukaryotic orders plotted against water temperature. Note that Vannellida (**A**, Amoebozoa), Cryomonadida (**B**, Cercozoa), and not further determined Trebouxiophyceae (**C**) decreased with increasing temperature, while Dactylopodida (**D**, Amoebozoa) increased.
